# Research Brief: Translation Initiation Represents an Acute Myeloid Leukemia Cell Vulnerability That Can Be Co-Targeted With BCL-2 Inhibition

**DOI:** 10.1101/2024.12.20.629809

**Authors:** Yoke Seng Lee, Jonathan D. Good, Noha Awais, Benjamin K. Eschle, Maia S. Lineberry, Jessica L. Teo, Sarah Bibeau, Morgan Wollman, George Karadimov, Nurefsan Sariipek, Jean Acosta, Jenny Noel, David Cucchi, Erica A.K. DePasquale, Prafulla C. Gokhale, Fadi J. Najm, Richard M. Stone, Jacqueline S. Garcia, Andrew A. Lane, Peter van Galen

**Affiliations:** Brigham and Women’s Hospital; Harvard Medical School; Ludwig Center at Harvard; Dana-Farber Cancer Institute; Boston University; Boston College; Sattler College; Amsterdam University Medical Center; Broad Institute of MIT and Harvard

## Abstract

Targeted therapies like Venetoclax have increased the options available to acute myeloid leukemia (AML) patients, but survival remains poor due to drug resistance and disease relapse. We found that the translation initiation factor *EIF4A1*, which unwinds complex mRNA structures in the 5’ UTR of oncogenic transcripts, is highly expressed in AML stem- and progenitor-like cells. Inhibiting eIF4A with the small molecule Zotatifin reduces translation of transcripts related to the cell cycle and survival. This results in downregulation of AKT, STAT-5, and MCL-1 and underlies synergy of Zotatifin with Venetoclax. The drug combination promotes apoptosis across AML genotypes, while the effect on healthy blood cells is limited. Using *in vivo* relapsed and refractory AML patient-derived xenograft models, the combination significantly suppressed tumor burden and prolonged survival of xenografted mice. These results support eIF4A-mediated protein translation as a therapeutic target in AML.

**SIGNIFICANCE:** Despite advances in targeted therapies, the 5-year survival rate for acute myeloid leukemia remains around 32%. The efficacy of existing treatments may be improved with the addition of Zotatifin, an inhibitor targeting the translation initiation factor eIF4A.

## INTRODUCTION

AML is an aggressive hematological malignancy arising from uncontrolled proliferation of myeloid progenitor cells in the bone marrow (BM) and blood. Standard AML treatment entails chemotherapy and allogeneic stem cell transplantation. However, refractory disease and high relapse rates remain major clinical problems, resulting in a 5-year survival of only 32%. Targeted therapies focused on patient-specific mutations have emerged as a new therapeutic approach, exemplified by FLT3 and isocitrate dehydrogenase-1/2 (IDH1/2) inhibitors. Other targeted therapies are directed against aberrant biological pathways, such as BCL-2 inhibitors that target dysregulated mitochondrial-driven apoptosis. The selective BCL-2 inhibitor Venetoclax coupled with hypomethylating agents such as Azacitidine (Ven/Aza) is now widely used for patients who are ineligible for intensive chemotherapy and in the relapsed/refractory setting (1,2). Nevertheless, the persistence of malignant cell populations that survive treatment underlies frequent disease recurrence (3). Comprehensive eradication of the malignancy is complicated by the heterogeneity of AML cells with regard to genomic landscapes and cellular composition (4–7). For example, Venetoclax resistance has been linked to myeloid-biased differentiation states and RAS mutations that upregulate anti-apoptotic MCL-1 (8,9). Other modes of resistance include the acquisition of escape mutations in BCL-2 which abrogate Venetoclax binding (10) and changes in mitochondrial structural integrity (11). Due to such resistance mechanisms, the response to targeted therapies is often short-lived.

Protein translation is a highly regulated process that cancer cells adapt to promote their proliferation and survival (12,13). Among the main phases of eukaryotic translation – initiation, elongation, and termination – initiation is the rate-limiting step. Translation initiation is mediated by the eukaryotic initiation factor 4F (eIF4F) complex, comprising the scaffold protein eIF4G, the cap-binding protein eIF4E, and the RNA helicase eIF4A (14). Mechanistically, eIF4A acts by unwinding complex structures in the 5’ UTR of select mRNAs, thereby promoting the translation of oncogenic transcripts (15,16). eIF4A has been implicated as a therapeutic target in cancer, and its inhibition by rocaglates like silvestrol or rocaglamide A has demonstrated preclinical efficacy with limited toxicity in healthy cells (17–21). eIF4A is highly expressed in leukemia relative to other cancer types, and the eIF4A inhibitor CR-131-B was found to synergize with Venetoclax in killing MOLM-14 AML cells (22,23). Recently, an eIF4A inhibitor with improved physicochemical properties, Zotatifin, was developed (24). Zotatifin is being evaluated in a phase 1/2 trial across multiple solid tumors and has shown favorable safety profiles (NCT04092673) (25,26). Zotatifin inhibits eIF4A by forming a stable ternary complex with eIF4A and its recognition motifs in the 5’ UTR of transcripts, thereby limiting the translation of oncogenic transcripts. Recently, Zotatifin was demonstrated to decrease BCL-2, BCL-XL and MCL-1 in lung cancer and synergize with RAS inhibitors to inhibit tumor growth both *in vitro* and *in vivo* (27). The evaluation of Zotatifin in hematological malignancies remains limited, with only one report in B-cell malignancies (28). Further, the combination of Zotatifin with BCL-2 inhibition has not been explored.

In this study, we identify eIF4A as a promising therapeutic target enriched in AML stem- and progenitor-like cell states, and find that its inhibition by Zotatifin is applicable to AML across diverse genotypes and differentiation states. By ribosome profiling, we show that Zotatifin decreases the translation efficiency of transcripts relating to PI3K/AKT/MTOR signaling and cell cycle pathways. Further, Zotatifin downregulates metabolic pathways and increases apoptosis and inflammation. By immunoblotting, Zotatifin decreases expression of MCL-1 at 4 hours and of AKT and STAT-5 at 24 hours, concomitant with BAK upregulation. We further demonstrate that the combination of Zotatifin and Venetoclax synergistically kills AML cells whilst sparing healthy human BM cells, indicating a therapeutic window for selective eradication of tumor cells. Finally, the combination achieved therapeutic efficacy in two clinically relevant AML patient-derived xenograft (PDX) models *in vivo* by curtailing tumor growth and prolonging survival. Taken together, our findings point towards a new therapeutic approach for treating AML patients.

## RESULTS

### *EIF4A1* is highly expressed and can be targeted in primitive AML cells

Leukemia cells exhibit increased protein synthesis compared to healthy hematopoietic stem cells (HSCs), resulting in dependence on protein translation elongation and termination pathways that can be therapeutically targeted (29,30). We hypothesized that protein translation initiation factors may be upregulated in leukemia cells. In single-cell gene expression data of healthy BM and AML cells, we found that the eIF4F complex member *EIF4A1* was upregulated in primitive leukemia cells (**Figure 1A, Supplementary Figure 1**) (6), which is supported by previous work that evaluated eIF4A expression in bulk primary AML cells (22).

**Figure 1.**
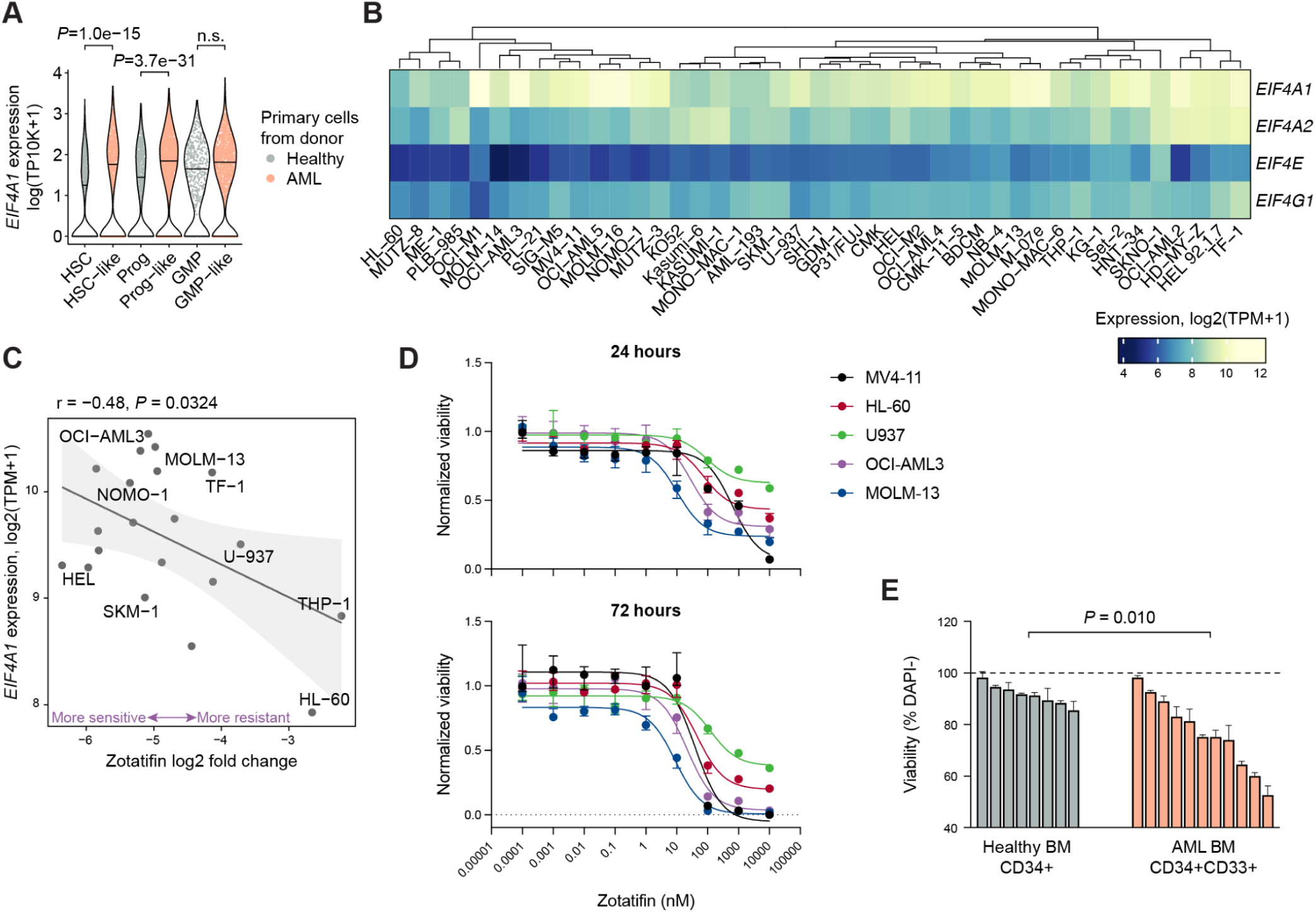
*EIF4A1* is highly expressed and can be targeted in primitive AML cell states. **A,** Sina plot shows expression of *EIF4A1* in individual cells analyzed by single-cell RNA-sequencing on 5 healthy and 16 AML BM samples (6). *P*-values were calculated using the Wilcoxon test with Bonferroni correction. **B,** Heatmap shows expression of eIF4F complex members across 43 AML cell lines analyzed by bulk RNA-seq (31). **C,** Scatter plot shows the reduction in AML cell viability after Zotatifin treatment (2.5 μM, five days, x-axis) and *EIF4A1* expression (y-axis) (32). Higher expression is associated with lower cell survival. **D,** Line graphs show half-maximal inhibitory concentration (IC50) curves for five AML cell lines treated for 24 or 72 hours with serial 10-fold titrated doses of Zotatifin from 0.1 pM to 10 μM. Results are representative of 2-4 independent experiments. Error bars represent mean±SD of n=3 technical replicates. **E,** Bar plot shows the percent viability after treating primary BM cells from healthy donors and AML patients with Zotatifin (100 nM, 24 hours). Viability of flow cytometry-gated CD34+ HSPCs from healthy BM samples and aberrant CD34+CD33+ cells from AML BM samples was normalized to DMSO treatment (dotted line at 100%). Each bar represents a biological replicate (n = 8 healthy donors, n = 11 for AML patients) and data is shown as mean ± SD of n = 3 technical replicates. *P*-value was calculated by unpaired t-test. HSC: hematopoietic stem cell, Prog: progenitor, GMP: granulocyte-macrophage progenitor, HSPC: hematopoietic stem and progenitor cells, BM: bone marrow.

Mining DepMap bulk RNA-seq data showed that *EIF4A1* is also highly expressed across 43 AML cell lines, and integration with PRISM drug treatment data indicated a correlation between *EIF4A1* expression and sensitivity to the eIF4A inhibitor Zotatifin (**Figure 1B-C**) (31). The PRISM drug studies were performed with Zotatifin at 2.5 µM for five days (32). To determine the extent to which it is effective at lower doses and shorter exposures, we treated 5 AML cell lines with titrated doses of Zotatifin for 24 hours and 72 hours and assessed viability using Cell-TiterGlo assay. Zotatifin reduced viability in all five cell lines, with a mean IC50 of 113.1 nM (range 11.0-399.0 nM) at 24 hours and 63.8 nM (range 12.8-138.6 nM) at 72 hours, suggesting applicability against AML across diverse genotypes and differentiation states (**Figure 1D** and **Supplementary Table 1**).

Finally, to test the potential presence of a therapeutic window, we cultured healthy BM cells and AML cells in StemSpan SFEM-II medium that supports hematopoietic cell/leukemic cell growth and treated the cells with Zotatifin (100 nM). We compared the viability of CD34+ hematopoietic stem and progenitor cells (HSPCs) from healthy BM and CD34+CD33+ cells from AML BM. This aberrant immunophenotype, which was largely absent in healthy BM but comprised 12.3% to 67.1% of CD45+ cells in AML samples, was used as a potential proxy for AML stem and progenitor cells. After 24 hours of treatment, we found increased cell death in Zotatifin-treated AML cells compared to healthy BM cells by flow cytometry (*P*=0.010, **Figure 1E**). These results suggest that AML cell survival selectively depends on the activity of eIF4A.

### Zotatifin downregulates AKT, metabolic, and cell cycle pathways, and upregulates inflammatory and apoptotic pathways

Next, we sought to determine how Zotatifin treatment affects gene expression and protein translation. To address this, we performed RNA-seq to measure gene expression and Ribo-seq to measure ribosome-protected fragments (RPFs, **Figure 2A**) (33). We treated MOLM-13 cells with DMSO (control) or Zotatifin (100 nM) for 4 hours, lysed the cells, isolated total RNA and ribosome-protected mRNA, and performed sequencing. Following alignment, we first assessed the position of RPFs along transcripts. We observed increased ribosome occupancy in the 5’ UTR and around the start codon, but not around the stop codon (**Figure 2B, Supplementary Figure 2A**). This is consistent with Zotatifin-induced immobilization of the eIF4F complex at the beginning of transcripts (34). We evaluated the RNA-seq data to identify genes with altered expression and Ribo-seq to identify genes with altered ribosome occupancy in Zotatifin compared to DMSO conditions, using DE-Seq2 to analyze both datasets separately (35). To determine pathways and biological processes that are affected by Zotatifin, we evaluated Hallmark gene sets using gene set enrichment analysis (GSEA). Gene sets that were downregulated by Zotatifin transcriptionally (based on RNA-seq) and translationally (Ribo-seq data) were related to metabolism, cell cycle and proliferation, and signaling pathways (**Figure 2C, Supplementary Figure 2B-D, Supplementary Table 3**). Downregulated metabolic processes include fatty acid metabolism, oxidative phosphorylation, and genes encoding components of the peroxisome. Downregulated cell cycle and proliferation processes include MYC and E2F targets; while downregulated signaling pathways include PI3K/AKT/MTOR and MTORC1 signaling. Gene sets that were upregulated by Zotatifin in both RNA-seq and Ribo-seq were enriched with stress and immune responses including apoptosis, inflammation, and interferons. In addition, upregulated signaling pathways include TNFα, TGFβ, and TP53 pathways.

**Figure 2.**
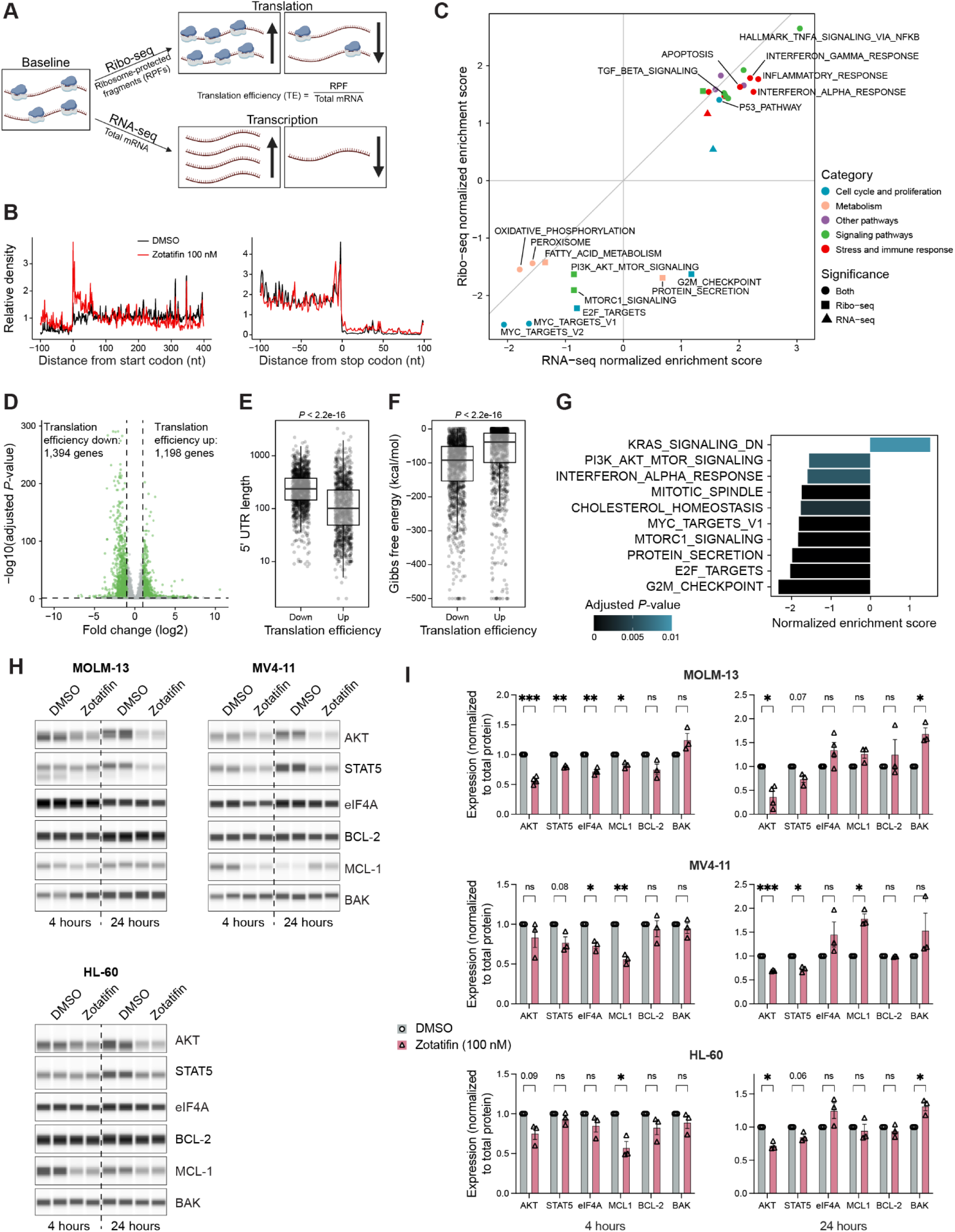
Zotatifin downregulates AKT, metabolic and cell cycle pathways, upregulates inflammatory and apoptotic pathways, and decreases STAT-5, and MCL-1 *in vitro*. **A,** Diagram illustrates the principles of ribosome profiling. Ribo-seq captures ribosome-protected mRNA fragments (RPFs), while RNA-seq measures the whole transcriptome. The ratio of RPFs to total mRNA is used to calculate translation efficiency (TE). Created with Biorender. **B,** Metagene plots for the relative densities of RPFs across all transcripts following 4-hour treatment of MOLM-13 cells with control (DMSO, black) or Zotatifin at 100 nM (red). X-axis represents the distance from the start codon (left) and stop codon (right). **C,** Correlation plot shows the normalized enrichment scores of RNA-seq (x-axis) and Ribo-seq (y-axis) following GSEA (Hallmark pathways), with selected biological processes annotated and color-coded. Shape indicates significance (*P*<0.05). For a full list, refer to **Supplementary Table 3**. **D,** Volcano plot shows the number of transcripts with upregulated or downregulated translation efficiency (TE) following Zotatifin treatment (*P*<0.05, 2-fold change). **E and F,** Box plots show the (**E**) 5’ UTR length and (**F**) 5’ UTR complexity (more negative Gibbs free energy means more complex 5’ UTR) of transcripts, categorized by transcripts with significantly increased or decreased TE. **G,** Bar plot shows normalized enrichment scores of Hallmark pathways following GSEA based on changes in TE following Zotatifin treatment. **H and I,** Western blot analysis of target protein expression following 4 or 24 hours treatment of MOLM-13, MV4-11 and HL-60 cells with 100 nM Zotatifin. **H,** Representative images of 3-4 independent experiments. Each experimental condition was performed with technical duplicates. **I,** Quantified protein expression across 3-4 independent experiments as in (H), normalized to total protein and DMSO control. Data represent mean±SEM across n=3-4 independent experiments each with technical duplicates. * *P*<0.05, ** *P*<0.01, *** *P*<0.001, **** *P*<0.0001, ns not significant, *P*-values between 0.05 and 0.1 are indicated. Welch’s t-test.

Collectively, we found that Zotatifin decreases processes and pathways relating to metabolism, cell survival, and proliferation, and increases stress/immune responses. Considering that Zotatifin’s target eIF4A promotes protein translation (12), we determined the translation efficiency of every transcript by calculating the ratio of RPFs to total mRNA counts within each gene’s coding sequence using DeltaTE (36). Zotatifin decreased the translation efficiency of 1,394 genes and increased the translation efficiency of 1,198 genes (**Figure 2D**). Transcripts with decreased translation efficiency harbored significantly longer and more complex 5’ UTR structures as indicated by a more negative Gibbs free energy (**Figure 2E-F**). To determine if Zotatifin-induced translation efficiency alterations affect specific gene sets, we performed GSEA. This revealed that Zotatifin decreased the translation efficiency of transcripts in the PI3K/AKT/MTOR signaling and cell cycle pathways (MYC, E2F targets, G2M checkpoint, **Figure 2G**), supporting that eIF4A inhibition by Zotatifin causes translational downregulation of oncogenic pathways.

We next performed western blotting to measure proteins that promote AML cell survival and growth. We treated MOLM-13, MV4-11, and HL-60 cells with 100 nM Zotatifin for 4 hours and 24 hours, and assessed key regulators of oncogenic signaling pathways and mitochondrial apoptosis. Representative images are shown in **Figure 2H** and total protein-normalized levels from at least three separate experiments are shown in **Figure 2I**. Total AKT was downregulated in all three cell lines at the 24-hour time point. STAT-5, representing JAK/STAT signaling, was also downregulated albeit to a lesser extent. We assessed the anti-apoptotic BCL-2 family proteins BCL-2 and MCL-1. MCL-1 was downregulated in all three cell lines at the 4-hour time point, which aligns with its short half-life (37). Both MCL-1 and eIF4A initially decreased but rebounded or increased after 24 hours, highlighting a dynamic response to Zotatifin. There were no major changes to BCL-2 expression. Consistent with apoptosis, Zotatifin increased expression of BAK in MOLM-13 and HL-60 cells after 24 hours (**Figure 2I**). Overall, Zotatifin decreases oncogenic drivers AKT, STAT-5 and eIF4A, decreases anti-apoptotic MCL-1, and increases pro-apoptotic BAK.

### Zotatifin synergizes with Venetoclax in AML cell lines

Because Zotatifin decreased AKT and MCL1 expression, which have been implicated in resistance to the BCL-2 inhibitor Venetoclax (38–42), we sought to determine if combining Zotatifin with Venetoclax could be synergistic. To this end, we treated MOLM-13, MV4-11, HL-60 and OCI-AML3 cell lines for 24 hours with titrated doses of Zotatifin and Venetoclax. These cell lines represent a range of AML differentiation states and genotypes, including *FLT3*, *TP53*, and *NRAS* mutations (**Supplementary Table 1**). While either drug alone elicited a dose-dependent decrease in AML cell viability, the combination elicited greater efficacy (**Supplementary Figure 3**). Applying synergy calculations using SynergyFinder, which employs the zero interaction potency (ZIP) model to quantify synergy by comparing observed combination responses to expected responses, demonstrated significant synergy between Zotatifin and Venetoclax across all four AML cell lines (ZIP synergy scores ranging from 29.7 to 57.2, **Figure 3A**). In a representative example, 50 nM of either drug alone resulted in a viability of 68.6-84.4%, but both drugs together decreased viability to 0.97% in MV4-11 cells (**Figure 3B**). For HL-60 cells, which are relatively primitive and sensitive to Venetoclax, we observed little further killing when combining Zotatifin with Venetoclax at 100 nM; however, strong synergy was still observed when using both drugs at 50 nM.

**Figure 3.**
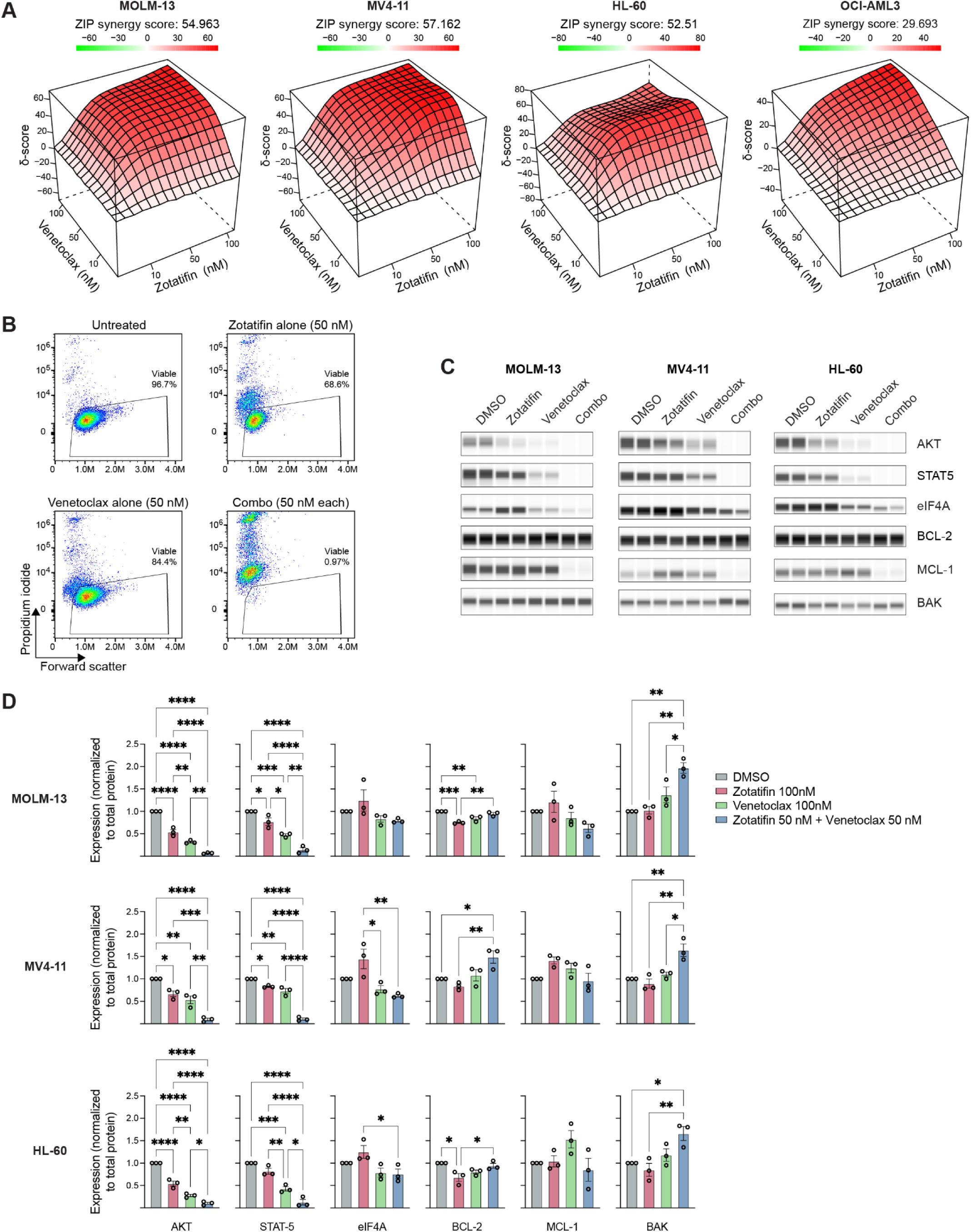
Zotatifin and Venetoclax synergize to kill AML cell lines. **A,** Synergy plots generated using SynergyFinder 3.0 for MOLM-13, MV4-11, HL-60 and OCI-AML3 cells treated for 24 hours with 10 nM, 50 nM, or 100 nM of Zotatifin or Venetoclax, alone or in combination, with DMSO as control. Representative plots of at least 2 independent experiments are shown. **B,** Representative flow cytometry plots show % viable MV4-11 cells following 24-hour treatment of DMSO control, Zotatifin alone (50 nM), Venetoclax alone (50 nM), or their combination (50 nM each). **C and D,** Western blot analysis of target protein expression following 24 hours treatment of MOLM-13, MV4-11 and HL-60 cells with 100 nM Zotatifin, 100 nM Venetoclax, or their combination at 50 nM each. (**C**) Representative images of 3 independent experiments. Each experimental condition was performed with technical duplicates. (**D**) Quantified protein expression across 3 independent experiments as in (C), normalized to total protein and DMSO control. Data represent mean±SEM across n=3 independent experiments each with technical duplicates. * *P*<0.05, ** *P*<0.01, *** *P*<0.001, **** *P*<0.0001, one-way ANOVA.

To dissect the mechanistic basis of this synergy, we treated MOLM-13, MV4-11, and HL-60 cells for 24 hours with Zotatifin or Venetoclax at 100 nM, or their combination at 50 nM each as determined to be synergistic. Zotatifin alone decreased AKT and STAT-5, as did Venetoclax alone, consistent with previous findings (43,44). Both proteins were further decreased by the combination (**Figure 3C** and **D**). Increased MCL-1 in MV4-11 cells treated with Zotatifin alone was similar to the 24-hour time point shown in **Figure 2G-H**, and a decrease was appreciable with the combination, but these changes did not reach statistical significance based on total protein-normalized values (**Figure 3C** and **D**). BAK was increased by the combination compared to monotherapies and DMSO control, consistent with apoptosis (**Figure 3B** and **C**). Overall, the combination reduced oncogenic AKT and STAT-5 and enhanced pro-apoptotic BAK expression.

### Zotatifin synergizes with Venetoclax against primary AML cells *in vitro* and *in vivo*

To test the clinical translatability of our findings, we wanted to determine if AML cells are selectively sensitive to Zotatifin plus Venetoclax compared to healthy cells. We cultured primary BM cells from healthy individuals (n=13) and AML patients (n=11, blasts 67.3±4.7% mean±sem) and treated them for 24 hours with either drug alone or their combination (**Supplementary Table 2**). We quantified cell viability by flow cytometry on total CD45+ leukocytes. Within this limited time frame, the combination treatment decreased viability in healthy BM to 87.5% compared to 71.2% in AML (**Figure 4A**). To assess if the combination had a different effect on diverse cell types, we further gated on CD34+ HSPCs from healthy BM and CD34+CD33+ putative AML stem and progenitor cells from AML BM, as well as major cell populations including CD33+ myeloid, CD19+ B, CD3+ T, and CD56+ NK cells (**Supplementary Figure 4A**). In AML samples, the combination decreased the viability of CD33+ myeloid cells to 65.6% and CD34+CD33+ cells to 49.9%, compared to 95.3% in healthy BM CD33+ myeloid cells and 80.9% in healthy BM CD34+ HSPCs (**Figure 4A**), highlighting a potential therapeutic window. In both healthy BM samples and AML samples, the effect of Zotatifin and Venetoclax on CD19+ B cells, CD3+ T cells, and CD56+ NK cells was limited, as the average viability measures all remained above 76%, further supporting limited toxicity in normal blood cells (**Figure 4A**).

**Figure 4.**
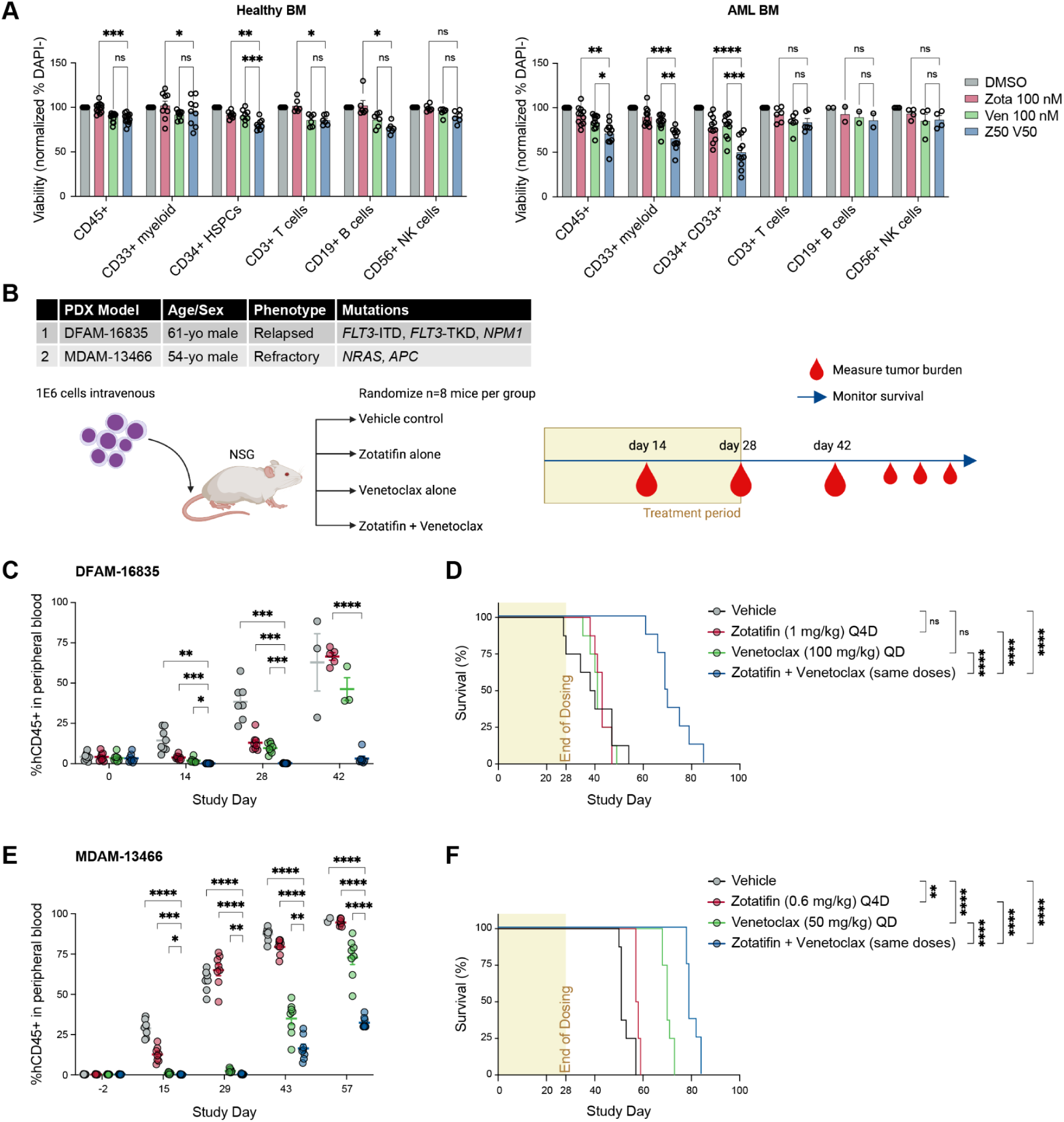
Zotatifin and Venetoclax synergize to kill primary AML cells *in vitro* and *in vivo*. **A,** Viability of healthy control (n=13) and AML bulk BM cells (n=11) treated for 24 hours with Zotatifin 100 nM, Venetoclax 100 nM, or their combination at 50 nM each (Z50 V50). Data normalized to DMSO control. Significance is shown for comparisons between combination and monotherapies only. The data for healthy CD34+ and AML CD34+CD33+ cells with Zotatifin monotherapy are also in Figure 1E and repeated for completeness. Some missing data points are due to cell types that did not reach a minimum of 300 cells to quantify viability (e.g., several AML samples were devoid of T, B and NK cells). **B,** Schematic of experimental design. Immunodeficient NSG mice (n=8 per group) were engrafted intravenously with 1 million AML cells from the indicated PDX models and treated for 28 days with Zotatifin intraperitoneally every 4 days (Q4D), Venetoclax once daily (QD) by oral gavage, or their combination at the indicated doses. Tumor burden was measured as the % human CD45+ cells in mouse peripheral blood every 14 days. Created with Biorender. **C,** Scatter plot shows tumor burden across study days by flow cytometry in DFAM-16835. Significance is only shown for comparisons between combination and other groups. **D,** Kaplan-Meier plot shows mouse survival curves across groups for DFAM-16835. **E,** Scatter plot shows tumor burden across study days in MDAM-13466. Significance is only shown for comparisons between combination and other groups. **F,** Kaplan-Meier plot shows mouse survival curves across groups for MDAM-13466. * *P*<0.05, ** *P*<0.01, *** *P*<0.001, **** *P*<0.0001; two-way ANOVA in (A), (C) and (E); log-rank test in (D) and (F).

The gold standard in preclinical drug efficacy studies is to test PDX models. We first determined the tolerability profile of Zotatifin and Venetoclax in immunodeficient Nod.Cg-Prkdc^scid^IL2rg^tm1Wjl^/SzJ (NSG) mice (n=4 per group). We treated these mice with drug doses that have been used in other studies (41,45) and demonstrated that a combination of Zotatifin at 1 mg/kg every four days and Venetoclax at 100 mg/kg daily for 14 days was tolerable in these animals, without significant impact on body weight (**Supplementary Figure 4B**).

Next, we evaluated the *in vivo* efficacy of Zotatifin and Venetoclax using one relapsed (DFAM-16835) and one refractory (MDAM-13466) AML PDX model, reflecting clinical scenarios with a critical need for new therapies (**Figure 4B** and **Supplementary Table 2**). In the first model (DFAM-16835), we engrafted mice with AML cells with complex cytogenetics from a 61-year-old patient at relapse following chemotherapy and allogeneic stem cell transplant. When peripheral blood measurement of human CD45+ cells reached a mean of 3.8±2.5%, we initiated treatment with monotherapies or the combination for 28 days (n=8 mice per group), in line with 28-day treatment cycles with Venetoclax-based regimens in clinical practice (46). We monitored tumor burden at 14-day intervals by measuring the % human CD45+ cells in the peripheral blood (**Figure 4B**). At the end of treatment (day 28), tumor burden was significantly decreased by the combination (mean 0.25%, range 0.10-0.50%) as compared to vehicle control (mean 38.4%, range 23.3-57.5%), Zotatifin monotherapy (mean 13.0%, range 8.5-24.1%), and Venetoclax monotherapy groups (mean 9.7%, range 4.7-13.7%, all *P*<0.001, **Figure 4C**). Strikingly, the median survival was 39 days for the vehicle control group, compared to 70 days for the Zotatifin + Venetoclax group (*P*<0.0001) (**Figure 4D**). Neither monotherapy was sufficient to confer a significant survival benefit in this aggressive PDX model, supporting the synergistic activity of eIF4A and BCL-2 inhibition.

In the second model (MDAM-13466), we engrafted mice with AML cells harboring *NRAS* and *APC* mutations from a 54-year-old patient with refractory disease following Decitabine treatment (**Figure 4B** and **Supplementary Table 2**). When peripheral blood measurement of human CD45+ cells reached a mean of 0.4±0.2% (MDAM-13466), we initiated treatment. To assess the robustness of the response and maximize clinical translatability, we lowered the dose of Zotatifin to 0.6 mg/kg and Venetoclax to 50 mg/kg, maintaining the same dose frequency and route of administration. Across all time points of tumor burden measurement, both on-treatment and post-treatment, the combination significantly decreased the percentage of human CD45+ cells relative to all other treatment groups (**Figure 4E**). For example, on study day 43, tumor burden remained significantly lower following the combination (mean 16.5%, range 7.4-28.8%) as compared to vehicle control (mean 87.3%, range 80.0-92.4%, *P*<0.0001), Zotatifin monotherapy (mean 79.4%, range 70.4-83.7%, *P*<0.0001), and Venetoclax monotherapy groups (mean 35.0%, range 15.7-47.9%, *P*<0.01, **Figure 4E**). Accordingly, the combination significantly extended survival relative to all other treatment groups (**Figure 4F**). Median survival was 51 days for the vehicle control group, compared to 79 days for the Zotatifin + Venetoclax group (*P*<0.0001) (**Figure 4F**). Despite the lower dose, Zotatifin or Venetoclax monotherapy was sufficient to confer a significant survival benefit in this model. Together, these *in vivo* data support the translational potential of combining Zotatifin and Venetoclax in clinically challenging cases of AML.

## DISCUSSION

In this study, we found eIF4A to be enriched in AML, particularly in primitive cell states relative to their healthy counterparts, underscoring its potential as a therapeutic target. Inhibition of eIF4A by Zotatifin decreases AML cell viability across genetically diverse AML cell lines and primary cells, suggesting that Zotatifin may be broadly applicable in AML. Using ribosome profiling, we found that Zotatifin decreases metabolic and cell cycle/proliferation pathways, including PI3K/AKT/MTOR signaling, and increases stress signaling and inflammatory pathways, including TP53, TNFα, and interferons. At the protein level, Zotatifin decreased AKT, STAT-5, and MCL-1, which have been associated with Venetoclax resistance, and was found to synergize with Venetoclax to kill AML cells both *in vitro* and *in vivo*. This lays the groundwork to develop Zotatifin and its combination with Venetoclax for the treatment of AML.

The potential of targeting eIF4A has been investigated in various cancers due to its importance for oncogene translation and relative safety in healthy tissues, supported by tolerability of Zotatifin in a phase 1/2 trial for solid tumors (25,26). This therapeutic window has previously been attributed to cancer cell addiction to elevated protein translation to maintain oncogenic signaling, which is consistent with our findings (12,18,19,47). Indeed, targeting translation elongation (29) and termination (48) has shown promising results in myelodysplastic syndromes (MDS) and AML. Recent work by Fooks and colleagues revealed that eIF4A inhibition of AML cells via CR-131-B perturbs the metabolic status of cells by decreasing mitochondrial membrane potential and the rate of ATP synthesis via glycolysis and oxidative phosphorylation (23). By ribosome profiling, we also found that Zotatifin perturbs metabolic pathways, including downregulation of oxidative phosphorylation and fatty acid metabolism. Future experiments are warranted to investigate to what extent the role of eIF4A in survival depends on the cellular context (differentiation states and genetics). In addition, different eIF4A inhibitors can exhibit partially overlapping yet diverse mechanisms of action, and Zotatifin may differ from earlier inhibitors in its specific mechanism of protein translation inhibition. For example, a study across more than 200 rocaglates has found that different inhibitors exhibit diverse mRNA-targeting spectra, and that selectivity for polypurine sequences is not universally conserved across all eIF4A-targeting compounds (49). In addition, a recent study found that eIF4A can bind polypurine sequences in the coding sequence (CDS) and rocaglamide A can inhibit translation elongation in addition to blocking translation initiation (34). Supporting the idea that specific rocaglate-like eIF4A inhibitors exhibit diverse mechanisms, CR-131-B has been reported to decrease the expression of all anti-apoptotic BCL-2 family members BCL-2, BCL-XL, and MCL-1 (23), while in our hands Zotatifin only decreased MCL-1 expression after 4 hours, a finding that was conserved across multiple cell lines. There is evidence, however, that Zotatifin’s mechanism may be tumor-specific, as concerted decreases in BCL-2, BCL-XL, and MCL-1 have been reported in lung cancer models (27). Our findings that Zotatifin decreases proliferation pathways are consistent with decreases in MYC, CDK4 and cyclin D1 from an earlier report in B cell lymphoma both *in vitro* and *in vivo* (28). While our study establishes Zotatifin as a strong candidate for AML treatment, ongoing drug development may reveal further opportunities for eIF4A inhibition (50).

Analysis of the pathways and proteins that are altered by Zotatifin led us to test its synergy with Venetoclax. Reported Venetoclax resistance pathways include PI3K/AKT signaling and metabolic reprogramming through redox adaptation and alterations in amino acid, fatty acid, and nicotinamide metabolism (38,42,51–54). We found that Zotatifin downregulates metabolic pathways including fatty acid metabolism, oxidative phosphorylation, and peroxisome genes, and AKT levels were further decreased by the combination with Venetoclax. This could limit escape mechanisms in AML cells. Another level of synergy may come from targeting different cell states. While Venetoclax primarily targets primitive cells (55), our results show that Zotatifin downregulates AKT and downstream mTOR signaling which are important for the survival of more differentiated cells (7). Thus, Zotatifin may target and eliminate residual AML cells that are not efficiently cleared by Venetoclax alone. The upregulation of TP53 signaling by Zotatifin suggests another potential mechanism of synergy by increasing apoptotic priming (23,56). Finally, the intriguing upregulation of inflammatory pathway genes by Zotatifin warrants further investigation. Overall, the pathways modulated by Zotatifin likely disrupt AML cell survival through multiple mechanisms.

Our studies using primary cells are supportive of a therapeutic window. In a time frame of 24 hours, AML cells with myeloid and progenitor cell phenotypes were selectively killed while healthy cells were spared. Further supporting the translational potential of our findings, combining Venetoclax and Zotatifin to treat two different clinically challenging PDX models prolonged survival reminiscent of long-term clinical benefit. This survival benefit in combination-treated mice relative to vehicle control and monotherapy groups provides a rationale for further exploration of tumor cell sensitivity at disease recurrence and the survival benefit of continuous treatment. Given the advent of novel triplet therapies in AML under preclinical and clinical investigation, future work may also evaluate combining Zotatifin and Venetoclax with other therapies in relapsed/refractory AMLs—an important clinical challenge. Overall, our studies represent a significant step forward by contributing mechanistic insight and pre-clinical data that strongly support targeting eIF4A-mediated translation initiation in AML.

## ACKNOWLEDGEMENTS

Y.S.L. is a Career Development Program (CDP) Special Fellow of The Leukemia & Lymphoma Society. P.v.G. is supported by the Ludwig Center at Harvard, the National Institutes of Health (NIH) (R33CA278393), the Starr Cancer Consortium, the Edward P. Evans Foundation, the MPN Research Foundation, the Vera and Joseph Dresner Foundation, a Research Scholar Grant from the American Cancer Society (RSG-24-1318769-01-CDP), and the Brigham Research Institute. Research reported in this publication was supported by the Ted and Eileen Pasquarello Tissue Bank in Hematologic Malignancies.

## AUTHOR CONTRIBUTIONS

Y.S.L., J.D.G., N.A., B.K.E., M.S.L., J.L.T., S.B., M.W., G.K., and N.S. performed experiments. Y.S.L., J.D.G., N.A., B.K.E., and P.v.G. analyzed data. J.A., J.N., D.C., E.A.K.D., A.A.L., R.S., J.S.G., and F.J.N. contributed resources and reagents. Y.S.L., B.K.E., P.C.G., and P.v.G. designed the experiments. Y.S.L., N.A., and P.v.G. wrote the paper.

## METHODS

### Inhibitors

Zotatifin (HY-112163) and Venetoclax (HY-15531) were purchased from MedChemExpress.

### Cell culture

AML cell lines MOLM-13, MV4-11, HL-60, OCI-AML3, and U937 were purchased from DSMZ or ATCC and cultured in RPMI-1640 medium containing 10% fetal bovine serum (FBS). Cell line authentication was confirmed by short tandem repeat (STR) DNA fingerprinting every 6 months. Mycoplasma testing was routinely conducted once every 3 months and cultures were confirmed to be negative by PCR. Primary BM samples were cultured in StemSpan™ Serum-Free Expansion Medium II (SFEM II) (StemCell Technologies #09605) containing a cytokine cocktail and UM729 in accordance with the manufacturer’s recommendations.

### Primary samples

Healthy BM samples were obtained from the sternum of individuals undergoing cardiac surgery at the Brigham and Women’s Hospital following informed consent in accordance with the institutional review board (IRB) protocol 2020P004103 at Mass General Brigham. In addition, human BM mononuclear cells (MNCs) were purchased from StemCell Technologies (#70001.2), while human BM CD34+ purified hematopoietic stem and progenitor cells (HSPCs) were purchased from Lonza Bioscience (#2M-101B) or StemCell Technologies (#70002.3). AML BM samples were obtained from the Pasquarello Tissue Bank at the Dana-Farber Cancer Institute, following informed consent in accordance with the IRB protocol 01-206 and secondary use protocol 20-123.

### Gene expression analysis

Single-cell RNA-sequencing data from 16 AML and 5 healthy BM samples was analyzed for genes related to eukaryotic translation initiation (6). In malignant cells and their healthy counterparts, the expression of these genes was normalized using the Seurat function NormalizeData. For sina/violin plots of *EIF4A1*, the adjusted P-value was calculated using the Wilcoxon test with Bonferroni correction as implemented by the Seurat function FindMarkers (57). To generate a heatmap of translation initiation factors, candidate genes were selected from the literature (14). Data was scaled using the Seurat function ScaleData and for each cell type, the average of scaled expression values was visualized using the ComplexHeatmap package (58). These analyses were performed in R for statistical computing version 4.3.1.

### Cell viability assays

For determination of Zotatifin half-maximal inhibitory concentration (IC50) values, AML cell lines MV4-11, HL-60, U937, OCI-AML3 and MOLM-13 were seeded at 0.5x10^6^ cells per ml (200 µl) in triplicate wells of 96-well plates and treated for 24 hours or 72 hours with serial 10-fold titrated doses of Zotatifin starting with a top dose of 10 µM. Cell viability was assessed by CellTiter-Glo assay (Promega G7570) according to the manufacturer’s recommendations and luminescence was measured using the Synergy H1 microplate reader (Agilent BioTek). IC50 curves were plotted based on the non-linear fit of [Inhibitor] vs. response (three parameters) model using GraphPad Prism version 8.0. For drug synergy experiments, AML cell lines were seeded as described above and treated for 24 hours with titrated doses of Zotatifin or Venetoclax alone (10 nM, 50 nM, and 100 nM), or their combination in various combinations. Cell viability was assessed on a Cytek Aurora flow cytometer (Cytek Biosciences) after staining with DAPI or propidium iodide at 1:2000 dilution for 15 mins. Flow cytometry data was analyzed using FlowJo version 10.8.1. Synergy scores and plots were generated using SynergyFinder 3.0 by uploading data to the web application (https://synergyfinder.fimm.fi/) (59). We used the viability readout under default settings including outlier detection and LL4 for curve fitting. For primary healthy and AML BM samples or purified CD34+ HSPCs, samples were treated for 24 hours with Zotatifin or Venetoclax alone at 100 nM, or their combination at 50 nM each, and viability was assessed by DAPI staining and flow cytometry using an antibody panel comprising CD3-FITC (BD #555332), CD14-PE-Cy7 (Beckman Coulter A22331), CD19-BV785 (BioLegend #302240), CD33-PE (BD #347787), CD34-APC-eFluor780 (Invitrogen #47-0349-42), CD45-PerCP (BD #340665), and CD56-APC (BD #341026).

### Western blot

AML cell lines were treated with 100 nM Zotatifin for 4 hours or 24 hours in parallel with 0.01% DMSO as the negative control and cell lysates were freshly collected in RIPA lysis buffer (EMD Millipore 20-188) containing protease inhibitor cocktail (ThermoFisher 78430) following the manufacturer’s recommendations. Protein concentration was measured using Bradford assay (Biorad 5000006) and western blotting was performed using Bio-Techne ProteinSimple Jess^TM^ system (SM-W004) following manufacturer’s recommendations, in technical duplicates across at least 3 independent experiments. STAT-5 (Cat. 94205), AKT (Cat. 9272S), eIF4A (Cat. 2013S), MCL-1 (Cat. 5453T), BAK (Cat. 12105S) and GAPDH (Cat. 2118S) antibodies were purchased from Cell Signaling Technologies. BCL-2 (12789-1-AP) antibody was purchased from Proteintech. Quantification of protein expression was expressed relative to DMSO control, after normalization to total protein load. AML cell lines were also treated with 100 nM Zotatifin alone, 100 nM Venetoclax alone, or their combination at 50 nM each for 24 hours before cell lysis and western blotting as described above.

### In vivo study

*In vivo* experiments were performed at Dana-Farber Cancer Institute at the Experimental Therapeutics core, in accordance with Dana-Farber Institutional Animal Care and Use Committee (IACUC) protocol 04-111. The tolerability profile of Zotatifin and Venetoclax combination was determined in immunodeficient Nod.Cg-*Prkdc^scid^IL2rg^tm1Wjl^*/SzJ (NSG) mice (n=4 per group) in which animals were treated with (1) Zotatifin 1 mg/kg every 4 days intravenously and Venetoclax 50 mg/kg daily by oral gavage, (2) Zotatifin 1 mg/kg and Venetoclax 100 mg/kg, or (3) Zotatifin 3 mg/kg and Venetoclax 100 mg/kg for 14 days or until mice were euthanized for ethical reasons. Zotatifin was formulated as a solution in 5% dextrose, while Venetoclax was formulated with 10% ethanol, 60% Phosal 50PG and 30% PEG400. Body weight of mice was monitored daily. The *in vivo* efficacy study was performed in two independent PDX models. The first PDX (DFAM-16835) is a relapsed AML model following high-dose chemotherapy and allogeneic stem cell transplant with complex cytogenetics, *FLT3*-ITD, *FLT3*-TKD, and *NPM1* mutations. The second (MDAM-13466) is a refractory AML model following Decitabine treatment with mutations in *NRAS* and *APC* (refer to **Supplementary Table 2** for more information on both models). Both models are derived from the Public Repository of Xenografts (PRoXe) (60). Mice were engrafted with 1 million cells intravenously before reaching 8 weeks of age. For DFAM-16835, treatment groups were vehicle control, Zotatifin 1 mg/kg every 4 days intraperitoneally, Venetoclax 100 mg/kg daily by oral gavage, and their combination at the same doses and routes of administration (n=8 mice per group). For MDAM-13466, treatment groups were vehicle control, Zotatifin 0.6 mg/kg every 4 days intraperitoneally, Venetoclax 50 mg/kg daily by oral gavage, and their combination at the same doses and routes of administration (n=8 mice per group). The % human CD45+ cells in mouse peripheral blood was monitored by flow cytometry every 14 days until the mice reached endpoint. Survival was also monitored until the mice reached endpoint.

### Ribosome profiling

MOLM-13 cells were treated for 4 hours with 100 nM Zotatifin or 0.01% DMSO as a control. To prepare samples for ribosome sequencing, 5-10 million cells were pelleted at 500 g for 5 minutes, the supernatant was discarded, and pellets were gently mixed with room temperature culture medium containing cycloheximide (final concentration 0.1 mg/ml). Cells were pelleted again, supernatant discarded, and pellets were gently mixed with pre-cooled PBS containing 0.1 mg/ml cycloheximide. Cells were pelleted again, supernatant discarded, and pellets were snap-frozen in liquid nitrogen for ribosome sequencing. To prepare samples for RNA sequencing, 5-10 million cells were pelleted and washed with pre-cooled PBS. Cells were pelleted again and TRIzol was added at 1 ml per million cells and pipetted repeatedly until no cell clumps remained. Samples were then snap-frozen in liquid nitrogen and sent to CD Genomics for ribosome sequencing and RNA-sequencing (https://www.cd-genomics.com/ribosome-profiling.html).

### Identification of differential translation efficiency genes

The quantification of gene expression was performed with the assistance of CD Genomics services who provided Ribo-seq and RNA-seq read counts. These gene counts were normalized using the DESeq2 statistical model through size factor estimation, followed by model fitting to account for the condition, sequencing type, and their interaction. Translation efficiency for all genes was then assessed using the deltaTE (ΔTE) integrative analysis approach in R, where ΔTE was calculated as the ratio of RPFs to total mRNA counts (ΔTE=RPF/mRNA) (36). As part of the DESeq2 package, the empirical Bayes technique was used to calculate log fold changes, and a Benjamini-Hochberg (BH) procedure was performed to control for the false discovery rate (FDR). An adjusted *P*-value threshold of 0.05 was used to identify significant changes in translation efficiency with a minimum 2-fold change.

### Differential gene expression and pathway enrichment analysis

Differential gene expression analysis of RNA-seq and Ribo-seq data was performed using DESeq2 (v.1.40.2) in R. Using NCBI identifications from the MANE select dataset (GRCh38.v1.3.ensembl), the genes were filtered for canonical transcripts to resolve any duplicates. The log2FoldChanges for ΔTE, RNA-seq, and Ribo-seq were then sorted in descending order to use as input for pathway enrichment analysis. This analysis was conducted using fast gene set enrichment analysis (FGSEA v.1.28.0) in R with a threshold of Benjamini-Hochberg (BH)-adjusted *P*-value < 0.05 and 10,000 gene permutations (61). We considered the Hallmark pathways (MsigDB v.7.5.1). The results of FGSEA were derived as normalized enrichment scores and their respective *P*-values. To assess the relationship between mRNA abundance and protein synthesis in significantly enriched pathways (*P* < 0.05), we plotted the normalized enrichment scores for RNA-seq against Ribo-seq data.

### Statistical analysis

Statistical analysis was performed using GraphPad Prism 8.0. Normality of data was assessed by Shapiro-Wilk test. Comparisons between two groups were performed using paired or unpaired t-tests according to experimental design. For comparisons between three or more groups, one-way or two-way analysis of variance (ANOVA) with Bonferroni’s multiple comparisons test was used. To assess synergy and calculate zero interaction potency (ZIP) scores, we used SynergyFinder 3.0. ZIP scores exceeding 10 are strongly indicative of an interaction between two drugs being synergistic (59). The Kaplan-Meier method was used to generate survival curves, and log-rank test for comparison between groups. Results are considered statistically significant when the *P*-value is <0.05.

### Code availability

Analysis of translation initiation factors in healthy BM and AML cell subpopulations is shared on https://github.com/petervangalen/reanalyze-aml2019. Analysis of DepMap and PRISM data is shared on https://github.com/petervangalen/AML-DepMap-Insights-2024. Analysis of Ribo-Seq data is shared on https://github.com/petervangalen/ProteinTranslation.

## TABLE LEGENDS

**Supplementary Table 1. IC50 of Zotatifin at 24 hours and 72 hours across 5 AML cell lines.** Table shows the half maximal inhibitory concentration (IC50) of Zotatifin at 24 hours and 72 hours in MV4-11, MOLM-13, HL-60, U937, and OCI-AML3 cell lines, as well as notable genetic mutations and FAB classification of differentiation states. LOH: loss of heterozygosity. M: male, F: female, Dx: diagnosis, na: no data available.

**Supplementary Table 2. Patient demographics and genetic information.** Information for all primary AML samples and the AML PDX model used in this study.

**Supplementary Table 3. Gene set enrichment analysis of Zotatifin-treated MOLM-13 cells.** Normalized enrichment scores (NES), *P*-values, and leading edge genes from GSEA using Hallmark gene sets. MOLM-13 cells were treated with Zotatifin (100 nM) for 4 hours, followed by RNA-seq and Ribo-seq.

**Supplementary Figure 1.**
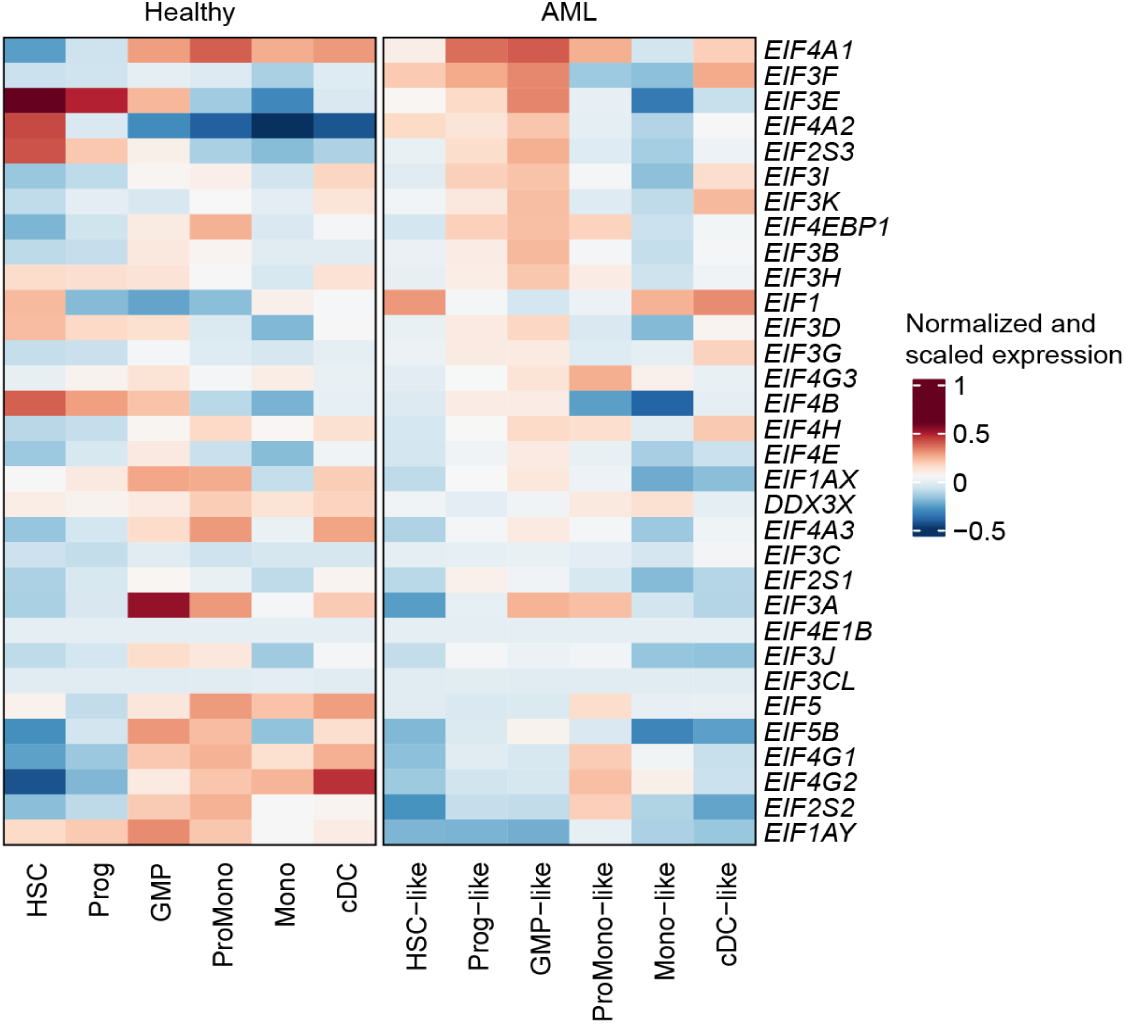
Expression of translation initiation factors in healthy and AML cell types by single-cell RNA-sequencing. Heatmap shows normalized expression of genes related to translation initiation (14) in indicated cell types. Heatmaps were generated using published single-cell RNA-sequencing data on 16 AML and 5 healthy BM samples (6). Rows are ordered by the gene with the highest expression in HSC-like, Prog-like, and GMP-like AML cells. HSC: hematopoietic stem cell, Prog: progenitor, GMP: granulocyte-macrophage progenitor, ProMono: promonocyte, Mono: monocyte, cDC: classical dendritic cell.

**Supplementary Figure 2.**
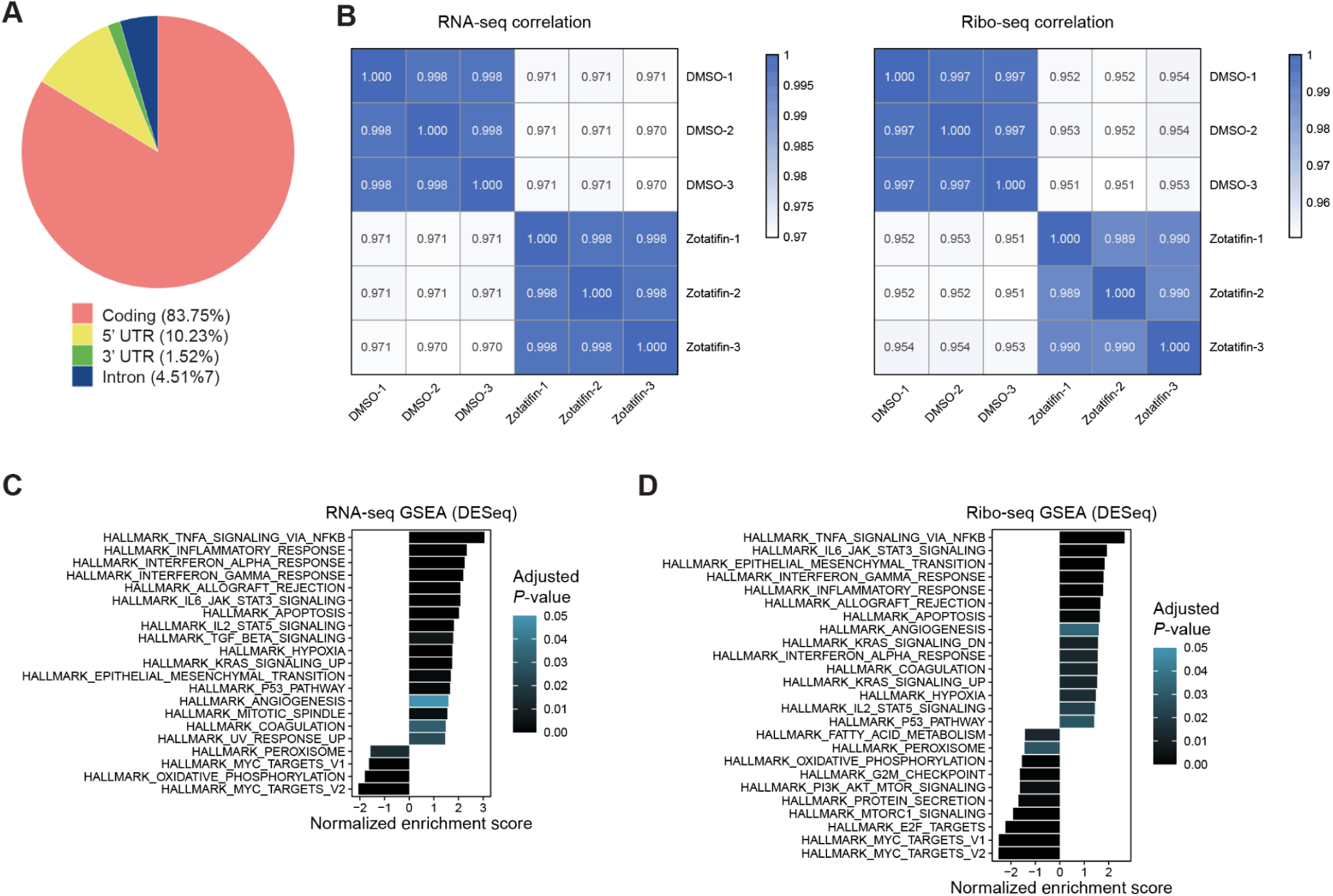
Ribosome profiling of MOLM-13 cells treated with Zotatifin. **A,** Pie chart shows the distribution of ribosome-protected fragments (RPFs) across the coding sequence (CDS), 5’ UTR, 3’ UTR, or intronic region of mRNAs assessed by ribosome profiling. A representative example of technical triplicates is shown for DMSO control. **B,** Heatmap shows correlation of gene counts between technical triplicates for paired RNA-seq (left) and Ribo-seq (right) of DMSO control and Zotatifin-treated MOLM-13 cells (4 hours). **C and D,** Bar plots show normalized enrichment scores of Hallmark pathways following GSEA of Zotatifin-treated MOLM-13 cells analyzed by (**C**) RNA-seq or (**D**) Ribo-seq.

**Supplementary Figure 3.**
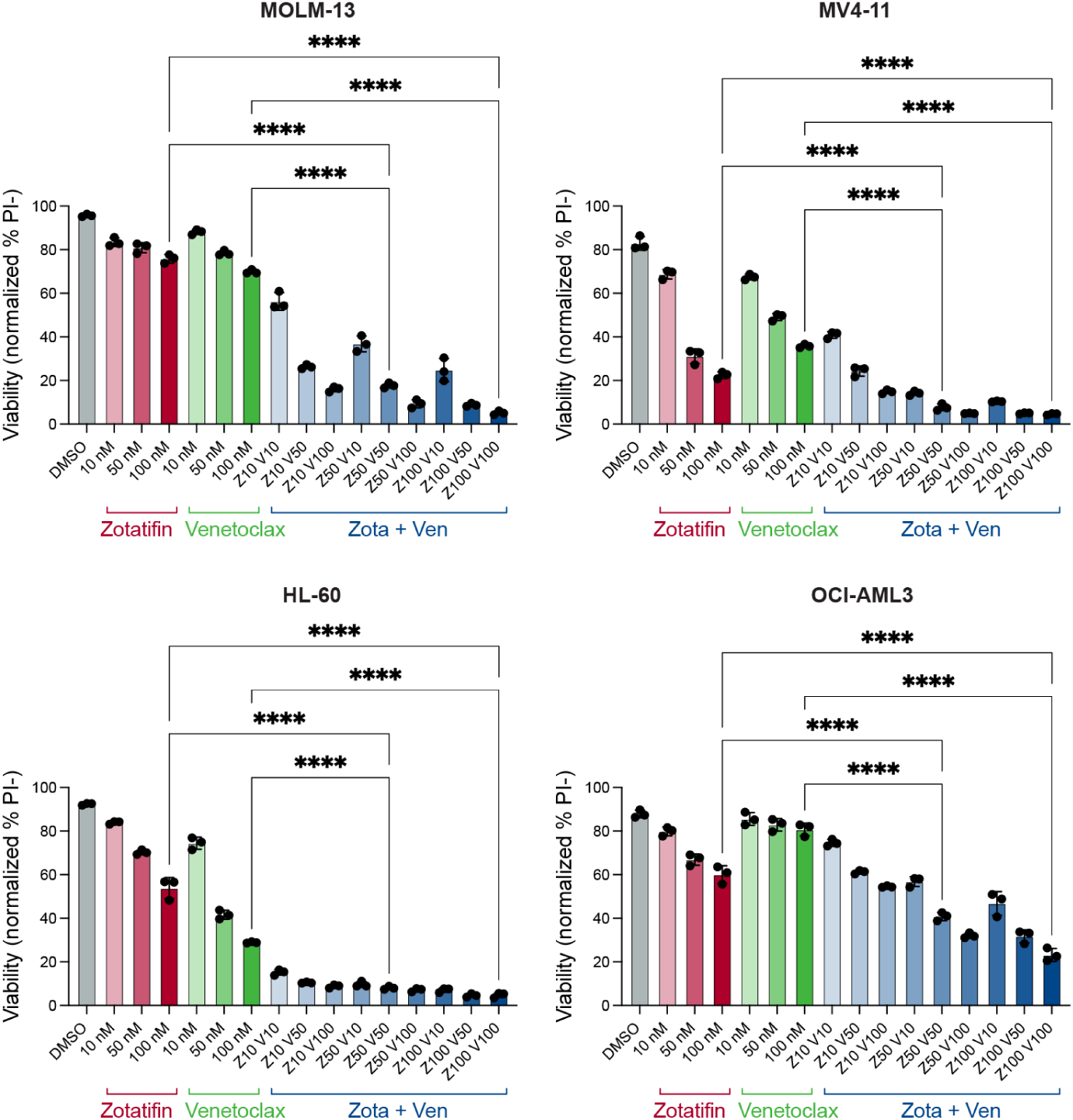
Zotatifin and Venetoclax synergize to kill AML cells *in vitro*. Bar plots show viability of MOLM-13, MV4-11, HL-60 and OCI-AML3 cells treated for 24 hours with indicated doses of Zotatifin or Venetoclax, alone or in combination. Viability was measured as the percentage of DAPI negative cells by flow cytometry. Representative plots of at least 2 independent experiments are shown. Data points represent technical triplicates within each experiment. Significance is only shown for comparisons between the top dose of Zotatifin or Venetoclax with their combination at 50 nM each or 100 nM each. **** *P*<0.0001; one-way ANOVA.

**Supplementary Figure 4.**
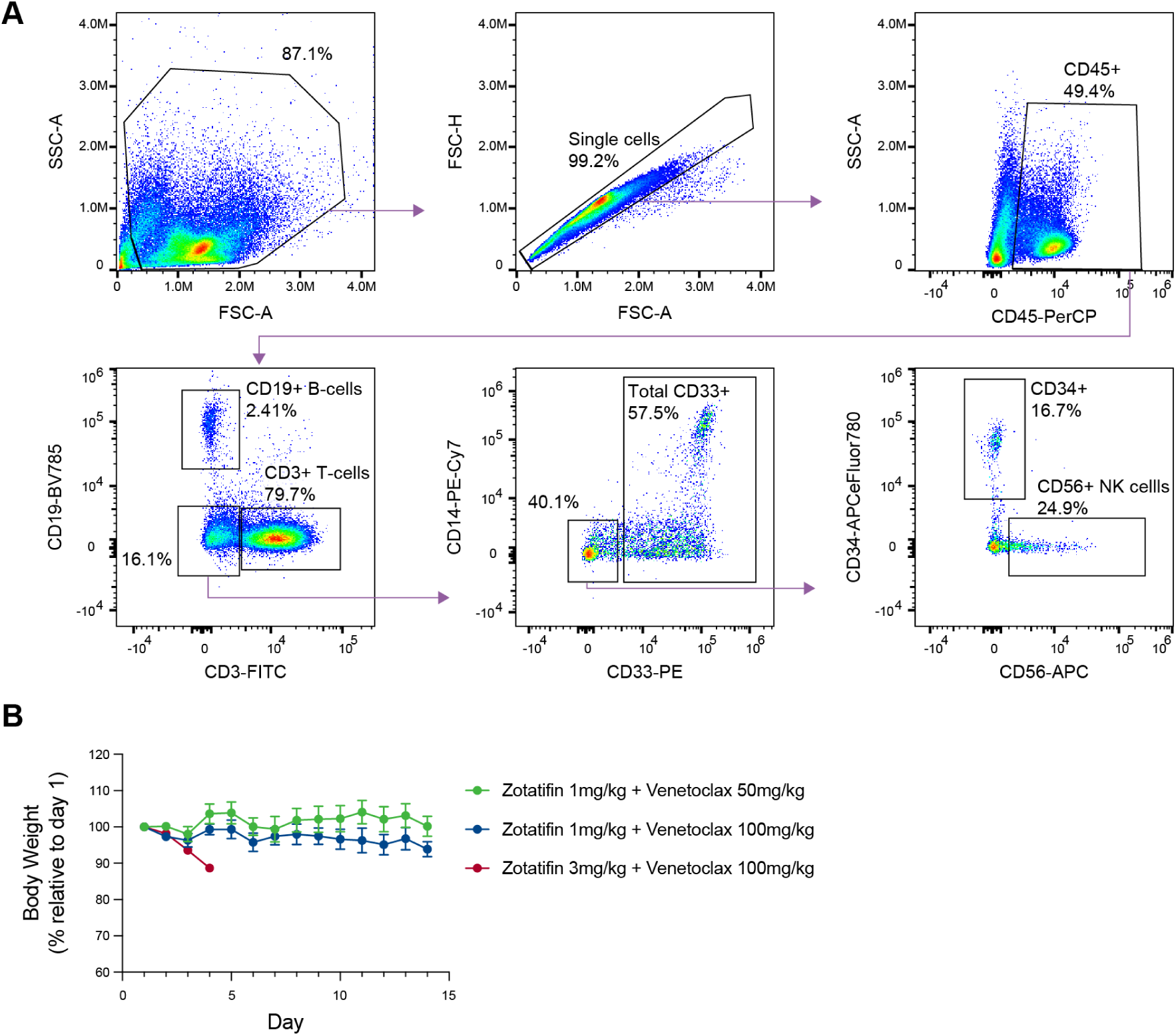
Gating strategy for primary human bone marrow and *in vivo* safety of Zotatifin and Venetoclax in NSG mice. **A,** Flow plots show gating strategy for a representative healthy BM sample delineating total CD45+ leukocytes, CD3+ T cells, CD19+ B cells, CD33+ myeloid cells, CD34+ HSPCs, and CD56+ NK cells. **B,** Line plots show *in vivo* safety study of Zotatifin and Venetoclax combination in NSG mice. Mice (n=4 per group) were treated with (1) Zotatifin 1 mg/kg every 4 days intravenously and Venetoclax 50 mg/kg daily by oral gavage, (2) Zotatifin 1 mg/kg and Venetoclax 100 mg/kg, or (3) Zotatifin 3 mg/kg and Venetoclax 100 mg/kg via the same routes of administration for 14 days or until mice were euthanized for ethical reasons. Body weight was measured daily and reported relative to that on day 1 of treatment.

## Notes

### Competing Interest Statement

The authors have declared no competing interest.

