## Supplementary Tables 1-3 for "Research Brief: Translation Initiation Represents an Acute Myeloid Leukemia Cell Vulnerability That Can Be Co-Targeted With BCL-2 Inhibition"

**Supplementary Table 1. IC50 of Zotatifin at 24 hours and 72 hours across 5 AML cell lines.**

| Cell line | Source | *FLT3* mutation | *RAS* mutation | TP53 mutation | Fusion gene | FAB status | IC50, 24h Zotatifin (nM) | IC50, 72h Zotatifin (nM) |
|---|---|---|---|---|---|---|---|---|
| MV4-11 | 10y/M/Dx | *FLT3*-ITD LOH | WT | WT | *MLL-AF4* | M5 | 399 | 43 |
| MOLM-13 | 20y/M/Relapse | *FLT3*-ITD (heterozygous) | WT | WT | *MLL-AF9* | M5a | 11 | 13 |
| HL-60 | 35y/F/Dx | WT | *NRAS* Q61L | p53 mutant | na | M2 | 60 | 103 |
| U937 | 37y/M/Refractory | WT | WT | p53 mutant | *CALM-AF10* | na | 67 | 139 |
| OCI-AML3 | 57y/M/Dx | WT | *NRAS* Q61L | WT | p53 WT | M4 | 28 | 21 |

LOH: loss of heterozygosity
M: male
F: female
Dx: diagnosis

**Supplementary Table 2. Patient demographics and genetic information.**

| 24-h liquid culture primary cells | | | | | | | |
|---|---|---|---|---|---|---|---|
| **Sample ID** | **Age** | **Sex** | **Mutations** | **Blasts** | **Dx/Rel** | **Cytogenetics** | **AML Diagnosis** |
| Pt1 | 78 | M | *FLT3-ITD, IDH-1* | 86% | Unknown | +13 | Unknown |
| Pt2.T1* | 24 | M | *CBL, NPM1* † | 78% | Relapse | 46,XY | AML with recurrent genetic abnormalties |
| Pt2.T2* | 25 | M | *CBL, NPM1* † | 62% | Relapse | 46,XY | AML with recurrent genetic abnormalties |
| Pt2.T3* | 25 | M | *CBL, NPM1* † | 55% | Relapse | 46,XY,t(1;7)(p3?2;q22),add(2)(q24),add(17)(q24)[3]/46,XY[cp17] | AML with recurrent genetic abnormalties |
| Pt3 | 62 | F | *FLT3, NPM1, TET2* † | 49% | Unknown | 46,XX | AML with recurrent genetic abnormalties |
| Pt4 | 63 | M | Not detected † | 75% | Relapse | 46,XY | AML, Not Otherwise Specified |
| Pt5 | 32 | F | *CEBPA, FLT3, NPM1* † | 84% | Diagnosis | 46,XX | AML with recurrent genetic abnormalties |
| Pt6 | 69 | M | *NPM1, WT1, CTCF, FLT3, NF1* † | 81% | Diagnosis | 46,XY | AML with recurrent genetic abnormalties |
| Pt7 | 58 | F | *DNMT3A, FLT3, NPM1* † | 75% | Diagnosis | 46,XX | AML with recurrent genetic abnormalties |
| Pt8 | 69 | M | Unknown | 56% | Unknown | Complex | Unknown |
| Pt9 | 60 | F | Not detected | 39% | Unknown | Complex | Unknown |

| PDX model | | | | | | | | |
|---|---|---|---|---|---|---|---|---|
| **Sample ID** | **Age** | **Sex** | **Mutations** | **Blasts** | **Dx/Rel** | **Treatment** | **Cytogenetics** | **AML Diagnosis** |
| DFAM-16835 | 61 | M | *FLT3-ITD, FLT3-TKD, NPM1* | 80% | Relapse | Induction chemotherapy, Consolidation HiDAC, Allogeneic HSCT, MEC | 47,X,-Y,del(6)(q15q21),+8,+14, del(15)(q12q15)[17]/ 47,idem,der(1)t(1;1)(p36.1q44)[2]/ 47,idem,t(1;9)(q23;q34)[1] | AML with recurrent genetic abnormalties; M4/M5 |
| MDAM-13466 | 54 | M | *NRAS, APC* | 44% | Refractory | Decitabine | 46,XY,del(3)(q21q25)[16]; 45,idem,-14,-17,+mar[1];46,XY[3] | AML, Not Otherwise Specified |

* Different time points from the same patient

† Source: Rapid Heme Panel (PMID: 27339098)

HiDAC: High-Dose Cytarabine

HSCT: Hematopoietic Stem Cell Transplant

MEC: Mitoxantrone, Etoposide, and Cytarabine

**Supplementary Table 3. Gene set enrichment analysis of Zotatifin-treated MOLM-13 cells.**

| pathway | pval_RNA | padj_RNA | log2err_RNA | ES_RNA | NES_RNA | size_RNA | leadingEdge_RNA | pval_Ribo | padj_Ribo | log2err_Ribo | ES_Ribo | NES_Ribo | size_Ribo | leadingEdge_Ribo | significance |
|---|---|---|---|---|---|---|---|---|---|---|---|---|---|---|---|
| HALLMARK_MYC_TARGETS_V2 | 0.0000 | 0.0001 | 0.5756 | -0.5566 | -2.0630 | 57 | UNG, SLC19A1, HK2, TN | 0.0000 | 0.0000 | 0.7477 | -0.6213 | -2.5058 | 57 | HK2, SLC19A1, HSPE1, | Both |
| HALLMARK_OXIDATIVE_PHOSPHORYLATION | 0.0000 | 0.0000 | 0.6273 | -0.3936 | -1.7908 | 198 | CYB5A, IDH1, CPT1A, A | 0.0001 | 0.0005 | 0.5188 | -0.3105 | -1.5463 | 199 | IDH1, VDAC3, CYB5A, N | Both |
| HALLMARK_MYC_TARGETS_V1 | 0.0001 | 0.0005 | 0.5384 | -0.3583 | -1.6300 | 199 | CAD, HSPE1, RRP9, VD | 0.0000 | 0.0000 | 1.0476 | -0.4998 | -2.4891 | 199 | HSPE1, CAD, GSPT1, X | Both |
| HALLMARK_PEROXISOME | 0.0053 | 0.0147 | 0.3183 | -0.3866 | -1.5699 | 93 | NR1I2, HSD11B2, IDH1, | 0.0129 | 0.0281 | 0.3108 | -0.3249 | -1.4392 | 93 | IDH1, HSD11B2, NR1I2, | Both |
| HALLMARK_FATTY_ACID_METABOLISM | 0.0259 | 0.0532 | 0.1462 | -0.3105 | -1.3528 | 145 | SLC22A5, IDH1, PPARA, | 0.0052 | 0.0131 | 0.4070 | -0.3010 | -1.4209 | 135 | SLC22A5, IDH1, HSPH1, | Ribo-seq |
| HALLMARK_ESTROGEN_RESPONSE_EARLY | 0.0266 | 0.0532 | 0.1462 | -0.2963 | -1.3176 | 168 | SLC22A5, ADCY1, TSKU | 0.0365 | 0.0652 | 0.0785 | 0.3481 | 1.3360 | 162 | PTGES, MSMB, DHRS2, | None |
| HALLMARK_APICAL_SURFACE | 0.1588 | 0.2647 | 0.0526 | -0.3701 | -1.2361 | 35 | SLC2A4, CD160, NTNG1 | 0.3782 | 0.4502 | 0.0418 | -0.2931 | -1.0420 | 32 | SLC2A4, TMEM8B, SCU | None |
| HALLMARK_WNT_BETA_CATENIN_SIGNALING | 0.2377 | 0.3714 | 0.0424 | -0.3351 | -1.1455 | 39 | JAG2, NKD1, FZD8, PSE | 0.2113 | 0.2709 | 0.0590 | -0.3172 | -1.1645 | 37 | JAG2, SKP2, AXIN2, PSE | None |
| HALLMARK_BILE_ACID_METABOLISM | 0.3605 | 0.5026 | 0.0354 | -0.2565 | -1.0417 | 93 | NR1I2, SLC29A1, KLF1, | 0.1621 | 0.2383 | 0.0830 | -0.2652 | -1.1541 | 84 | SLC29A1, IDH1, CROT, | None |
| HALLMARK_APICAL_JUNCTION | 0.3540 | 0.5026 | 0.0375 | -0.2331 | -1.0365 | 168 | NRXN2, AMIGO1, NRTN | 0.4100 | 0.4659 | 0.0191 | 0.2712 | 1.0322 | 148 | SLIT2, ADRA1B, SDC3, | None |
| HALLMARK_GLYCOLYSIS | 0.5341 | 0.6540 | 0.0296 | -0.2162 | -0.9722 | 183 | CYB5A, CHST4, EFNA3, | 0.0575 | 0.0992 | 0.1986 | -0.2446 | -1.2013 | 182 | PDK3, HK2, CHST4, IDH | None |
| HALLMARK_PI3K_AKT_MTOR_SIGNALING | 0.7913 | 0.9201 | 0.0213 | -0.2097 | -0.8539 | 95 | MAP2K6, SFN, PLA2G12 | 0.0010 | 0.0035 | 0.4551 | -0.3681 | -1.6306 | 93 | MAP2K6, SLA, SLC2A1, | Ribo-seq |
| HALLMARK_MTORC1_SIGNALING | 0.8797 | 0.9562 | 0.0215 | -0.1875 | -0.8531 | 198 | UNG, HK2, IDH1, POLR | 0.0000 | 0.0000 | 0.7050 | -0.3829 | -1.9066 | 198 | HK2, IDH1, SLC37A4, H | Ribo-seq |
| HALLMARK_E2F_TARGETS | 0.9613 | 1.0000 | 0.0202 | -0.1762 | -0.8014 | 200 | UNG, RPA1, DSCC1, TR | 0.0000 | 0.0000 | 0.8987 | -0.4462 | -2.2224 | 199 | GSPT1, XPO1, RBBP7, | Ribo-seq |
| HALLMARK_DNA_REPAIR | 1.0000 | 1.0000 | 0.0189 | -0.1371 | -0.5986 | 149 | PDE6G, NPR2, CDA, CC | 0.9971 | 1.0000 | 0.0053 | 0.1610 | 0.6127 | 148 | BOLA2, BCAM, EIF1B, D | None |
| HALLMARK_PROTEIN_SECRETION | 0.9818 | 1.0000 | 0.0119 | 0.1773 | 0.6775 | 93 | ABCA1, DOP1A, SH3GL | 0.0004 | 0.0014 | 0.4985 | -0.3828 | -1.6915 | 90 | ATP1A1, PAM, CLTC, SE | Ribo-seq |
| HALLMARK_REACTIVE_OXYGEN_SPECIES_PAT | 0.9289 | 0.9882 | 0.0133 | 0.2062 | 0.7028 | 49 | GPX3, IPCEF1, JUNB, F | 0.3902 | 0.4537 | 0.0449 | -0.2633 | -1.0303 | 48 | OXSR1, GSR, HMOX2, | None |
| HALLMARK_NOTCH_SIGNALING | 0.8698 | 0.9562 | 0.0151 | 0.2369 | 0.7215 | 29 | LFNG, HES1, DTX2, MA | 0.7249 | 0.7711 | 0.0139 | 0.2785 | 0.8285 | 30 | NOTCH3, HES1, LFNG, | None |
| HALLMARK_ADIPOGENESIS | 0.8219 | 0.9340 | 0.0136 | 0.2025 | 0.8536 | 187 | PPARG, GPX3, CAVIN1, | 0.0641 | 0.1068 | 0.1881 | -0.2416 | -1.1864 | 182 | SLC19A1, IDH1, SLC1A5 | None |
| HALLMARK_UNFOLDED_PROTEIN_RESPONSE | 0.5514 | 0.6564 | 0.0202 | 0.2434 | 0.9551 | 111 | TUBB2A, ATF3, CCL2, I | 0.0744 | 0.1200 | 0.1371 | -0.2675 | -1.2204 | 111 | ALDH18A1, ERO1A, HY | None |
| HALLMARK_ANDROGEN_RESPONSE | 0.5363 | 0.6540 | 0.0208 | 0.2520 | 0.9592 | 91 | SGK1, STEAP4, H1-0, P | 0.1532 | 0.2322 | 0.0867 | -0.2619 | -1.1575 | 90 | ABCC4, ALDH1A3, ADA | None |
| HALLMARK_HEME_METABOLISM | 0.5092 | 0.6528 | 0.0208 | 0.2340 | 0.9779 | 174 | BTG2, SLC6A8, H1-0, S | 0.8399 | 0.8749 | 0.0081 | 0.2127 | 0.8226 | 174 | BTG2, H1-0, DMTN, AC | None |
| HALLMARK_MYOGENESIS | 0.5068 | 0.6528 | 0.0210 | 0.2367 | 0.9780 | 159 | COL1A1, GPX3, SLC6A8 | 0.3626 | 0.4422 | 0.0210 | 0.2772 | 1.0575 | 152 | ACHE, PGAM2, FST, CC | None |
| HALLMARK_ESTROGEN_RESPONSE_LATE | 0.3796 | 0.5129 | 0.0256 | 0.2488 | 1.0316 | 164 | PTGES, DHRS2, SGK1, | 0.1165 | 0.1820 | 0.0423 | 0.3175 | 1.2186 | 162 | PTGES, DHRS2, AREG, | None |
| HALLMARK_XENOBIOTIC_METABOLISM | 0.3619 | 0.5026 | 0.0264 | 0.2504 | 1.0423 | 169 | PTGES, CYP1A1, AKR1 | 0.1826 | 0.2468 | 0.1041 | -0.2287 | -1.1057 | 163 | IDH1, ABHD6, SLC1A5, | None |
| HALLMARK_SPERMATOGENESIS | 0.3547 | 0.5026 | 0.0276 | 0.2731 | 1.0546 | 100 | SYCP1, H1-6, SHE, MAP | 0.5886 | 0.6540 | 0.0148 | 0.2619 | 0.9402 | 92 | H1-6, SYCP1, DPEP3, M | None |
| HALLMARK_COMPLEMENT | 0.2229 | 0.3596 | 0.0355 | 0.2689 | 1.1192 | 169 | OLR1, DPP4, ADRA2B, | 0.1818 | 0.2468 | 0.0326 | 0.3014 | 1.1627 | 169 | GZMB, OLR1, C1S, ADR | None |
| HALLMARK_G2M_CHECKPOINT | 0.1359 | 0.2342 | 0.0465 | 0.2771 | 1.1746 | 197 | BCL3, EGF, ARID4A, MT | 0.0001 | 0.0004 | 0.5384 | -0.3273 | -1.6274 | 196 | HSPA8, SQLE, SLC38A1 | Ribo-seq |
| HALLMARK_UV_RESPONSE_DN | 0.0547 | 0.0985 | 0.0768 | 0.3253 | 1.3086 | 131 | COL1A1, DUSP1, PPAR | 0.2037 | 0.2680 | 0.0312 | 0.3095 | 1.1561 | 126 | DUSP1, PPARG, DLC1, | None |
| HALLMARK_CHOLESTEROL_HOMEOSTASIS | 0.0551 | 0.0985 | 0.0785 | 0.3726 | 1.3589 | 70 | PPARG, TNFRSF12A, A | 0.2319 | 0.2898 | 0.0648 | -0.2676 | -1.1196 | 68 | PDK3, SQLE, ALCAM, F | None |
| HALLMARK_KRAS_SIGNALING_DN | 0.0349 | 0.0671 | 0.0972 | 0.3467 | 1.3750 | 119 | BTG2, PDCD1, SGK1, E | 0.0035 | 0.0111 | 0.2652 | 0.4250 | 1.5608 | 109 | BTG2, PDCD1, KRT1, S | Ribo-seq |
| HALLMARK_UV_RESPONSE_UP | 0.0097 | 0.0255 | 0.1849 | 0.3566 | 1.4547 | 147 | RRAD, BTG2, CYP1A1, | 0.1776 | 0.2468 | 0.0335 | 0.3093 | 1.1740 | 144 | RRAD, BTG2, LHX2, CY | RNA-seq |
| HALLMARK_COAGULATION | 0.0126 | 0.0314 | 0.1643 | 0.3769 | 1.4753 | 108 | THBS1, OLR1, DPP4, M | 0.0041 | 0.0122 | 0.2452 | 0.4226 | 1.5452 | 106 | THBS1, OLR1, C1S, TF | Both |
| HALLMARK_HEDGEHOG_SIGNALING | 0.0251 | 0.0532 | 0.1222 | 0.4987 | 1.5405 | 31 | HEY1, ACHE, SCG2, AD | 0.6939 | 0.7542 | 0.0146 | 0.2854 | 0.8492 | 30 | ACHE, HEY1, SCG2, VE | None |
| HALLMARK_MITOTIC_SPINDLE | 0.0012 | 0.0037 | 0.4551 | 0.3667 | 1.5509 | 193 | SYNPO, SHROOM1, CL | 1.0000 | 1.0000 | 0.0045 | 0.1419 | 0.5544 | 194 | PALLD, FARP1, CLIP2, | RNA-seq |
| HALLMARK_ANGIOGENESIS | 0.0207 | 0.0492 | 0.1353 | 0.5351 | 1.5904 | 26 | OLR1, VCAN, STC1, TH | 0.0171 | 0.0342 | 0.1306 | 0.5413 | 1.5875 | 28 | STC1, OLR1, APOH, VC | Both |
| HALLMARK_PANCREAS_BETA_CELLS | 0.0265 | 0.0532 | 0.1204 | 0.5986 | 1.6061 | 17 | MAFB, DPP4, FOXO1, N | 0.0357 | 0.0652 | 0.0925 | 0.5706 | 1.5170 | 18 | MAFB, NEUROD1, DPP4 | None |
| HALLMARK_P53_PATHWAY | 0.0003 | 0.0012 | 0.4985 | 0.3930 | 1.6565 | 188 | RRAD, JUN, IL1A, BTG2 | 0.0139 | 0.0289 | 0.1280 | 0.3617 | 1.4066 | 184 | RRAD, JUN, BTG2, IER | Both |
| HALLMARK_EPITHELIAL_MESENCHYMAL_TRAN | 0.0007 | 0.0022 | 0.4773 | 0.4087 | 1.6841 | 155 | THBS1, JUN, COL1A1, | 0.0000 | 0.0000 | 0.6436 | 0.4790 | 1.8292 | 153 | THBS1, GEM, JUN, TNC | Both |
| HALLMARK_KRAS_SIGNALING_UP | 0.0001 | 0.0004 | 0.5384 | 0.4191 | 1.7403 | 167 | IL1B, MAFB, FGF9, NR1 | 0.0044 | 0.0122 | 0.2321 | 0.3989 | 1.5241 | 154 | IL1B, SPARCL1, BIRC3, | Both |
| HALLMARK_HYPOXIA | 0.0000 | 0.0001 | 0.5756 | 0.4152 | 1.7472 | 185 | JUN, HAS1, DUSP1, HO | 0.0063 | 0.0150 | 0.1914 | 0.3799 | 1.4703 | 176 | JUN, HAS1, IER3, STC1 | Both |
| HALLMARK_TGF_BETA_SIGNALING | 0.0019 | 0.0057 | 0.4551 | 0.5194 | 1.7797 | 51 | THBS1, KLF10, SKIL, HI | 0.0266 | 0.0511 | 0.1004 | 0.4542 | 1.4804 | 49 | THBS1, PPP1R15A, KLF | RNA-seq |
| HALLMARK_IL2_STAT5_SIGNALING | 0.0000 | 0.0001 | 0.5933 | 0.4305 | 1.8106 | 182 | IRF4, PHLDA1, TNFRSF | 0.0094 | 0.0213 | 0.1563 | 0.3697 | 1.4324 | 178 | PHLDA1, TNFRSF18, IR | Both |
| HALLMARK_APOPTOSIS | 0.0000 | 0.0000 | 0.6901 | 0.4914 | 2.0156 | 151 | JUN, IL1A, IL1B, BTG2, | 0.0005 | 0.0017 | 0.4985 | 0.4275 | 1.6304 | 151 | JUN, IL1B, BTG2, IER3, | Both |
| HALLMARK_IL6_JAK_STAT3_SIGNALING | 0.0000 | 0.0000 | 0.6105 | 0.5650 | 2.0769 | 74 | JUN, IL1B, CCL7, CRLF | 0.0000 | 0.0001 | 0.5756 | 0.5534 | 1.9243 | 74 | JUN, IL1B, CRLF2, CCL | Both |
| HALLMARK_ALLOGRAFT_REJECTION | 0.0000 | 0.0000 | 0.7195 | 0.5002 | 2.0801 | 168 | IL11, IL1B, IRF4, CCL4, | 0.0001 | 0.0004 | 0.5384 | 0.4308 | 1.6577 | 166 | IL11, CCL4, ACHE, IL1B, | Both |
| HALLMARK_INTERFERON_GAMMA_RESPONSE | 0.0000 | 0.0000 | 0.8634 | 0.5196 | 2.1908 | 187 | IRF4, CCL7, FPR1, TNF | 0.0000 | 0.0000 | 0.6273 | 0.4587 | 1.7862 | 188 | IRF4, CCL7, C1S, OASL | Both |
| HALLMARK_INTERFERON_ALPHA_RESPONSE | 0.0000 | 0.0000 | 0.7749 | 0.5850 | 2.2482 | 96 | TXNIP, TMEM140, OASL | 0.0052 | 0.0131 | 0.2208 | 0.4278 | 1.5460 | 96 | C1S, OASL, CXCL10, CS | Both |
| HALLMARK_INFLAMMATORY_RESPONSE | 0.0000 | 0.0000 | 0.9326 | 0.5551 | 2.3316 | 181 | IL1A, IL1B, SGMS2, CCL | 0.0000 | 0.0000 | 0.6105 | 0.4602 | 1.7678 | 163 | IL1B, BTG2, CXCL8, CC | Both |
| HALLMARK_TNFA_SIGNALING_VIA_NFKB | 0.0000 | 0.0000 | 1.5698 | 0.7228 | 3.0502 | 189 | JUN, IL1A, MSC, IL1B, P | 0.0000 | 0.0000 | 1.3499 | 0.6806 | 2.6484 | 187 | CCL4, EGR2, GEM, JUN | Both |
